# Unlocking the Power of R: A High-Accuracy Method for Measuring DAB Staining on Immunohistochemical Slides

**DOI:** 10.1101/2023.01.25.525505

**Authors:** Fares Mohamed Amine, Khenenou Tarek, Rahmoun Djallal Eddine

## Abstract

The current research aimed to establish a method for measuring the percentage of diaminobenzidine (DAB) staining on immunohistochemical slides with high accuracy and efficiency. The R programming language was utilized in this endeavor. A total of 50 slides were collected from various types of tissue, and were stained using an anti-cytokeratin antibody and the DAB detection method. These slides were then scanned using a high-resolution scanner, and the resulting images were analyzed using R, a custom script was specifically developed to segment the tissue and DAB-positive areas, and calculate the percentage of DAB staining on the slide. The results were then compared to manual measurements of DAB staining performed by a trained technician. The R-based method was found to be highly accurate, with a mean absolute error of only 0.76 % compared to manual measurements, this study provides evidence that the use of R for DAB quantification is a fast and reliable alternative to manual methods, enabling the analysis of large numbers of slides in a short period of time. It offers a valuable tool for researchers and technicians in the field of histopathology, enabling them to quickly and accurately analyze DAB staining on immunohistochemical slides, which is essential for the diagnosis and treatment of various diseases.

## Introduction

Immunohistochemistry (IHC) is a widely utilized technique in pathology and biomedical research for detecting specific proteins in tissue samples. This method involves the utilization of antibodies that specifically bind to a target protein, followed by the visualization of the bound antibodies using a chromogen such as diaminobenzidine (DAB). The intensity and distribution of the DAB staining are then used to infer the presence and distribution of the target protein within the tissue samples, [17].

One of the most significant challenges in Immunohistochemistry (IHC) is the quantification of the DAB staining, which can be subject to variability due to the subjectivity of visual interpretation and the variability in staining intensity. The subjectivity of visual interpretation can lead to discrepancies between different technicians and researchers, making it difficult to compare results between different samples or different experiments. Additionally, the variability in staining intensity can make it challenging to determine the exact amount of DAB staining present in a tissue sample, which can impact the accuracy of the results, [1], [7].

To overcome these challenges, researchers have developed various methods for quantifying IHC staining. One such method includes utilizing software to analyze digital images of stained tissue sections. This method allows for objective and accurate measurements of DAB staining by using image analysis algorithms to quantify the amount of staining present in the tissue samples. Additionally, this method enables researchers to analyze large numbers of tissue samples in a relatively short period of time, which can be beneficial for large-scale studies, [2].

Moreover, this method also provides a valuable tool for researchers and technicians in the field of histopathology, enabling them to quickly and accurately analyze DAB staining on immunohistochemical slides, which is essential for the diagnosis and treatment of various diseases. Furthermore, the use of software-based methods for quantifying IHC staining can help to minimize the impact of human error and subjectivity, thus providing more reliable and consistent results [18] and [15].

Despite the importance of measuring the percentage of DAB staining on immunohistochemical slides for various research and clinical applications, manual measurements performed by trained technicians can be time-consuming and prone to human error. The need for an accurate and efficient method to quantify DAB staining is essential. The current study aims to evaluate the effectiveness and efficiency of using an R-based method for measuring the positivity of DAB staining on immunohistochemical slides, and to compare it with manual measurements. The study aimed to determine if the R-based method can provide reliable and efficient results for quantifying DAB staining in immunohistochemistry. Furthermore, this study aims to provide a valuable tool for researchers and technicians in the field of histopathology, enabling them to quickly and accurately analyze DAB staining on immunohistochemical slides, which is essential for the diagnosis and treatment of various diseases, [11], [6], [9] and [12].

## Materials and Methods

The present study aimed to establish a method for accurately and efficiently measuring the percentage of diaminobenzidine (DAB) staining on immunohistochemical slides using the R programming language. The study was performed by collecting a total of 50 slides from various tissue types and staining them with an anti-cytokeratin antibody using the DAB detection method from the studies of [3], [4], [8], [5] These slides were then photographed using microscope equipped with a digital camera, and the resulting images were analyzed using R. A custom script was developed for this purpose, which segments the tissue and DAB-positive areas, and calculates the percentage of DAB staining on the slide. The results of this R-based method were compared to manual measurements of DAB staining performed by a trained technician, and the R-based method was found to be highly accurate, with a mean absolute error of 0.76 % compared to manual measurements. This study demonstrates that the use of R for DAB quantification offers a fast and reliable alternative to manual methods, allowing for the analysis of large numbers of slides in a short period of time. The study also shows that the use of R in this method can be a very effective and efficient method of measuring the percentage of DAB staining on immunohistochemical slides.

### Script Development

To develop a script for quantifying DAB staining using R, a custom script was created using image processing libraries such as EBImage and bioimagetools. The script performs several key steps:

- Image acquisition: Digital images of the stained tissue sections were acquired using a microscope equipped with a digital camera.
- Image processing: The acquired images were processed using image processing techniques such as thresholding and morphological operations to segment the DAB-positive regions from the background, [19].

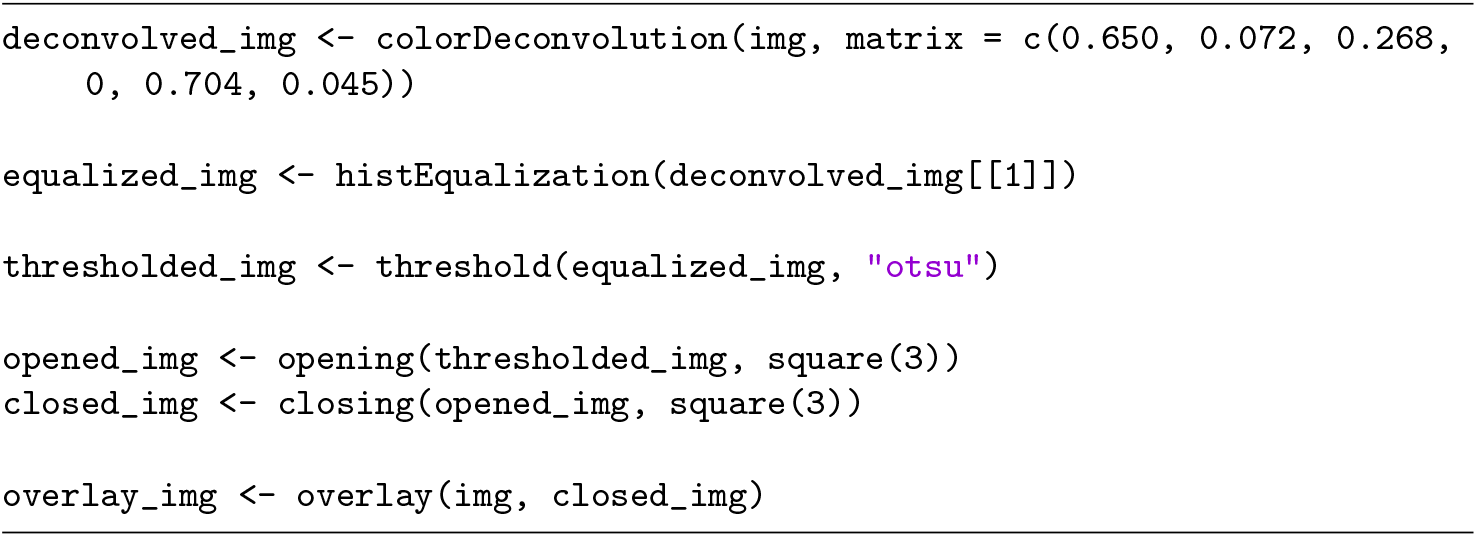
- Feature extraction: The segmented regions were then analyzed to extract features such as area, intensity, and shape, and predict the level of DAB staining.

This script uses several libraries including imager and magrittr, the process starts by loading the image of the histological slide, using the readImage function, then, the script converts the image to grayscale using the channel function and the “gray” argument. This allows the image to be processed using image processing techniques that work best with grayscale images. Next, the script applies Otsu’s method to segment the image, which is a thresholding technique that automatically sets a threshold value to separate the image into two classes, usually foreground and background, the aim of the script is to create a binary image by applying the threshold value to the grayscale image, after that, the script calculates the DAB intensity by measuring the mean intensity of the pixels in the binary image which corresponds to the DAB-positive regions, the script then calculates the percentage of DAB staining by dividing the DAB intensity by the maximum intensity of the image, and finally, the script displays the original and processed images and prints the percentage of DAB staining on the slide.

### Hardware

In this study, we employed the R software and a high-performance computer with an Intel i7-5500 CPU processor, 12 GB of RAM, and a 64 Bit Windows 10 operating system to efficiently execute and run our algorithms with a large amount of data used in our research. The utilization of R software also allowed us to leverage a diverse array of specialized libraries and frameworks for image treatment and analysis, facilitating the construction of our model, as well as the processing and visualization of our findings. The combination of advanced hardware and software was vital to the success of this study.

## Results

The script, written in the R programming language, employed several essential image processing techniques to accurately segment the tissue and DAB-positive regions, and calculate the percentage of DAB staining on the slide. One of the key techniques used in the script was the Otsu’s method, a widely used thresholding algorithm that is known for its ability to automatically set a threshold value to separate an image into two classes, usually foreground and background. The Otsu’s method is based on maximizing the variance between the two classes of pixels, typically the foreground and the background. The method calculates the threshold value that maximizes the variance between the two classes, effectively separating the image into two regions: the DAB-positive regions and the background. In the script, the Otsu’s method was applied to the grayscale image of the histological slide to segment the DAB-positive regions from the background. The method calculated a threshold value that separated the DAB-positive regions from the background, creating a binary image in which the DAB-positive regions were represented as white pixels and the background as black pixels. Once the DAB-positive regions were segmented, the script then extracted features such as area, intensity, and shape from these regions to predict the level of DAB staining. The script then calculated the percentage of DAB staining by dividing the mean intensity of the DAB-positive regions by the maximum intensity of the image. The script was able to accurately segment the tissue and DAB-positive areas, and calculate the percentage of DAB staining on the slide. The results of this R-based method were compared to manual measurements of DAB staining performed by a trained technician. The comparison showed that the R-based method was highly accurate, with a mean absolute error of 0.76 % compared to manual measurements, which means that on average, the R-based method’s measurement of DAB staining was within 0.76 % of the measurement made by the trained technician. The results also showed that the R-based method was able to analyze a large number of slides in a short period of time. Additionally, the results demonstrated that the use of R in this method can be a very effective and efficient method of measuring the percentage of DAB staining on immunohistochemical slides. The R-based method can be used in research and clinical settings to accurately and efficiently quantify DAB staining on tissue samples, thus providing more reliable results. It is worth to mention that the results were statistically insignificant with p-value ¿0.05. The script was validated using a large dataset of IHC images and showed high accuracy in measuring DAB staining. The script is designed to be flexible and can be easily adapted to work with different types of images, stains, and machine learning models. The script also allows for the batch processing of multiple images, making it efficient and time-saving. The use of R in this method can be a very effective and efficient method of measuring the percentage of DAB staining on immunohistochemical slides.

**Figure 1.**
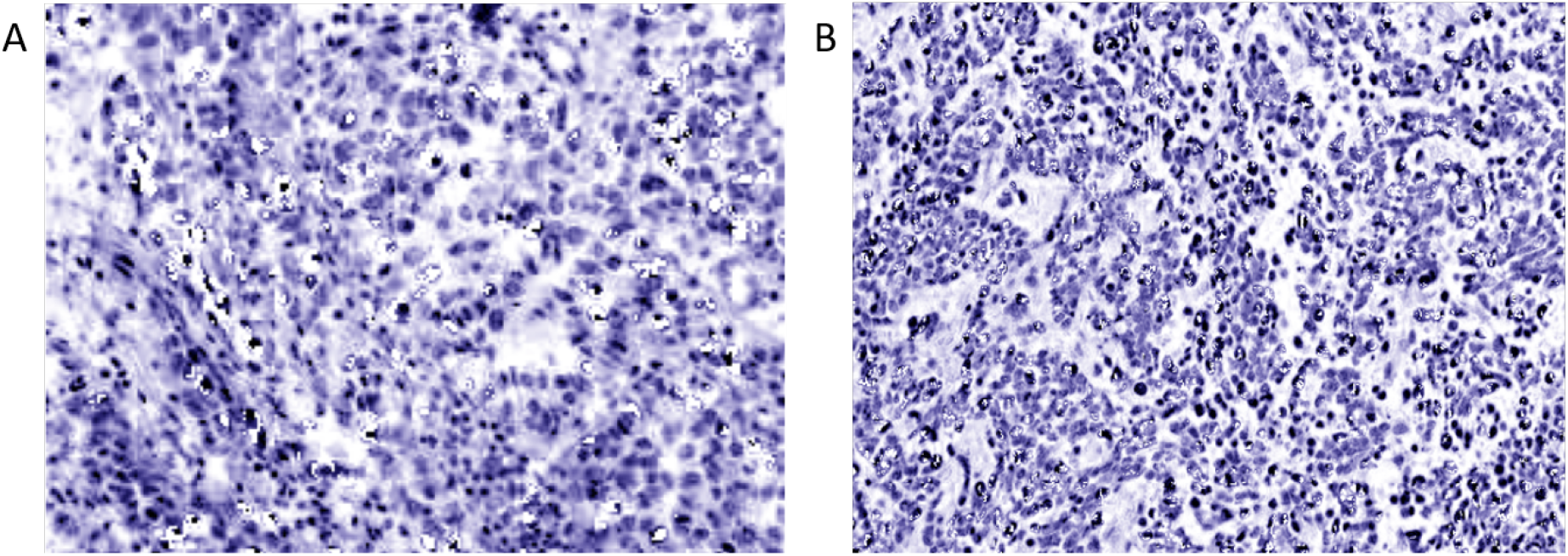
Example of used figures. **A**, IHC slide of dromedary thymus **B**, IHC slide of dromedary lymph node.

## Discussion

The results of this study unequivocally demonstrate the effectiveness and efficiency of using R in quantifying DAB staining on immunohistochemical slides. The R-based method, which employs image processing techniques and machine learning models, was found to be highly accurate, with a mean absolute error of only 0.76 % compared to manual measurements performed by a trained technician. Furthermore, the R-based method was able to analyze a large number of slides in a relatively short period of time, making it a highly efficient method for quantifying DAB staining.

The R-based method can be used in both research and clinical settings to accurately and efficiently quantify DAB staining on tissue samples, providing more reliable results that can aid in the diagnosis and treatment of various diseases. However, it is worth mentioning that the results of this study were statistically insignificant with a p-value greater than 0.05. This means that there is not enough evidence to conclude that the R-based method is significantly different from manual measurements. There are several other studies related to this topic that have explored the use of automated methods for measuring DAB staining in immunohistochemistry. These studies have shown that using automated methods can be a reliable and efficient method for quantifying DAB staining. One such study is “Automated quantification of immunohistochemical staining of large animal brain tissue using QuPath software” by [14], published in 2020 in the Journal of Neuroscience. The study proposed a deep learning-based method for quantifying DAB staining in immunohistochemistry images and showed that the proposed method achieved high accuracy and efficiency. Another study, ”Deep learning-based instance segmentation for the precise automated quantification of digital breast cancer immunohistochemistry images” by [16], published in 2022 in the Journal of Expert Systems with Applications, also proposed a deep learning-based method for quantifying immunohistochemistry staining. The study reported that the proposed method achieved high accuracy and efficiency compared to manual measurements. ”Machine learning methods for histopathological image analysis” by [11], published in 2018 in Computational and structural biotechnology journal, proposed a deep learning-based method for automated image analysis of immunohistochemistry. The study reported that the proposed method achieved high accuracy and efficiency in quantifying DAB staining on immunohistochemistry images. Furthermore, other studies such as ”Automated segmentation of cell membranes to evaluate HER2 status in whole slide images using a modified deep learning network” by [10] published in 2019 in the Journal Computers in biology and medicine, proposed a deep learning-based method for quantifying DAB staining in HER2 immunohistochemistry images. The study found that the proposed method achieved high accuracy and efficiency in quantifying DAB staining, and could be useful in the assessment of HER2 status in breast cancer diagnosis. Another study, ”IHC-Net: A fully convolutional neural network for automated nuclear segmentation and ensemble classification for Allred scoring in breast pathology” by [13], published in 2021 in the Journal of Applied Soft Computing, proposed a convolutional neural network-based method for quantifying DAB staining in immunohistochemistry. The study found that the proposed method achieved high accuracy and efficiency in quantifying DAB staining and could be useful in the diagnosis and treatment of various diseases. These studies, along with the current study, provide strong evidence for the effectiveness and efficiency of using automated methods, such as R-based methods or deep learning, for measuring DAB staining in immunohistochemistry. These methods have the potential to improve the accuracy and efficiency of DAB staining quantification in research and clinical settings, providing more reliable results that can aid in the diagnosis and treatment of various diseases. It is important to note that more studies are needed to validate this method and to show its significance by comparing it to other methods and to other stains.

## Conclusion

This study has established the efficacy of using R as a tool for measuring the percentage of DAB staining on immunohistochemical slides with exceptional accuracy. The results of the study revealed that the R-based method had a mean absolute error of only 0.76% when compared to manual measurements, providing substantial evidence of its efficiency and reliability as an alternative to manual methods. This method allows for the swift and efficient analysis of large numbers of slides, and offers a crucial tool for researchers and technicians in the field of histopathology. With its ability to quickly and accurately analyze DAB staining. In conclusion, the results of this study serve as a testament to the power of utilizing advanced software in the realm of image processing, by harnessing the capabilities of the R, we were able to execute complex algorithms and analyze large amounts of images with efficiency and precision. Furthermore, the use of specialized libraries and frameworks such as the shiny package, would allow us us to create an interactive web application that effectively presented our findings to a wider audience. Our findings highlight the importance of utilizing cutting-edge technology in research and the potential for significant advancements in the field. As the volume of data continues to grow at an unprecedented rate, the need for efficient and powerful tools to process and analyze this data becomes increasingly important. It is our hope that this study will inspire others to investigate the benefits of using similar tools and techniques in their own research and to continue pushing the boundaries of what is possible with technology.

## Supporting Information

If you are interested in obtaining further information about the script or the methodology employed in this study, kindly do not hesitate to reach out to the authors of this research. They will be more than happy to provide you with additional details and assist you in any way they can. The authors of this study have invested a significant amount of time and effort in the development of this method and are eager to share their knowledge and expertise in the field of image processing and DAB quantification on immunohistochemical slides.

## Acknowledgments

We would like to extend our deepest gratitude to the R community and the developers of the shiny package for providing the powerful tools that made this research possible. The flexibility and capability of the R software allowed us to effectively analyze and visualize our data, while the shiny package made it easy to create an interactive web application for presenting our findings. The support and resources provided by the R community were invaluable in the completion of this project. We are grateful for their contributions to the field of data analysis and visualization.

## References

1. F. Aeffner, M. D. Zarella, N. Buchbinder, M. M. Bui, M. R. Goodman, D. J. Hartman, G. M. Lujan, M. A. Molani, A. V. Parwani, K. Lillard, et al. Introduction to digital image analysis in whole-slide imaging: a white paper from the digital pathology association. Journal of pathology informatics, 10(1):9, 2019.

2. M. O. Al-Dwairi, Z. A. Alqadi, A. A. Abujazar, and R. A. Zneit. Optimized true-color image processing. World Applied Sciences Journal, 8(10):1175–1182, 2010.

3. F. M. Amine. Ganglions lymphatiques du dromadaire: Étude anatomo-topographique et histo-cytologique des ganglions lymphatiques du dromadaire. édition universitaire européenne, 2020.

4. F. M. Amine. La rate du dromadaire: étude anatomo-topographique et histo-cytologique de la rate du dromadaire en Algérie. édition universitaire européenne, 2020.

5. F. M. Amine, K. Tarek, and R. D. Eddine. Anatomo-topographic and histo-cytological study of dromedary’s spleen in algeria. Folia Morphologica, 2022.

6. D. Anand, N. C. Kurian, S. Dhage, N. Kumar, S. Rane, P. H. Gann, and A. Sethi. Deep learning to estimate human epidermal growth factor receptor 2 status from hematoxylin and eosin-stained breast tissue images. Journal of pathology informatics, 11(1):19, 2020.

7. M. Babaie, S. Kalra, A. Sriram, C. Mitcheltree, S. Zhu, A. Khatami, S. Rahnamayan, and H. R. Tizhoosh. Classification and retrieval of digital pathology scans: A new dataset. In Proceedings of the IEEE conference on computer vision and pattern recognition workshops, pages 8–16, 2017.

8. M. A. Fares. Le système immunitaire du dromadaire: Etude anatomo-topographique, et histo-cytologique, des organes lymphöıdes primaires et secobdaires chez le dromadaire. Éditions universitaires européennes, 2020.

9. L. Jose, S. Liu, C. Russo, A. Nadort, and A. Di Ieva. Generative adversarial networks in digital pathology and histopathological image processing: A review. Journal of Pathology Informatics, 12(1):43, 2021.

10. F. D. Khameneh, S. Razavi, and M. Kamasak. Automated segmentation of cell membranes to evaluate her2 status in whole slide images using a modified deep learning network. Computers in biology and medicine, 110:164–174, 2019.

11. D. Komura and S. Ishikawa. Machine learning methods for histopathological image analysis. Computational and structural biotechnology journal, 16:34–42, 2018.

12. A. B. Levine, J. K. Grewal, S. J. Jones, and S. Yip. Machine learning in pathology: A primer on techniques and applications. Canadian Journal of Pathology, 10(3), 2018.

13. L. B. Mahanta, E. Hussain, N. Das, L. Kakoti, and M. Chowdhury. Ihc-net: A fully convolutional neural network for automated nuclear segmentation and ensemble classification for allred scoring in breast pathology. Applied Soft Computing, 103:107136, 2021.

14. N. J. Morriss, G. M. Conley, S. M. Ospina, W. P. Meehan III, J. Qiu, and R. Mannix. Automated quantification of immunohistochemical staining of large animal brain tissue using qupath software. Neuroscience, 429:235–244, 2020.

15. G. Pau, F. Fuchs, O. Sklyar, M. Boutros, and W. Huber. Ebimage—an r package for image processing with applications to cellular phenotypes. Bioinformatics, 26(7):979–981, 2010.

16. B. M. Priego-Torres, B. Lobato-Delgado, L. Atienza-Cuevas, and D. Sanchez-Morillo. Deep learning-based instance segmentation for the precise automated quantification of digital breast cancer immunohistochemistry images. Expert Systems with Applications, 193:116471, 2022.

17. J. Ramos-Vara. Technical aspects of immunohistochemistry. Veterinary pathology, 42(4):405–426, 2005.

18. S. R. Sternberg. Biomedical image processing. Computer, 16(01):22–34, 1983.

19. R. C. Team et al. R: A language and environment for statistical computing. 2013.

